# Sugar feeding enhances gut immunity and protects against arboviral infection in the mosquito vector *Aedes aegypti*

**DOI:** 10.1101/2021.01.05.425375

**Authors:** Floriane Almire, Sandra Terry, Melanie McFarlane, Agnieszka M. Sziemel, Selim Terhzaz, Margus Varjak, Alma McDonald, Alain Kohl, Emilie Pondeville

## Abstract

As mosquito females require a blood meal to reproduce, they can act as vectors of numerous pathogens, such as arboviruses (*e.g.* Zika, dengue and chikungunya viruses), which constitute a substantial worldwide public health burden. In addition to blood meals, mosquito females can also take sugar meals to get carbohydrates for their energy reserves. It is now recognised that diet is a key regulator of health and disease outcome through interactions with the immune system. However, it has been mostly studied in humans and model organisms. So far, the impact of sugar feeding on mosquito immunity and in turn, how this could affect vector competence for arboviruses has not been analysed. Here, we show that sugar feeding increases and maintains antiviral immunity in the digestive tract of the main arbovirus vector *Aedes aegypti*. Our data demonstrate that the gut microbiota does not mediate the sugar-induced immunity but partly inhibits it. Importantly, sugar intake prior to an arbovirus-infected blood meal further protects females against infection with arboviruses from different families, highlighting a broad antiviral action of sugar. Sugar feeding blocks arbovirus initial infection and dissemination from the gut, lowers infection prevalence and intensity, thereby decreasing transmission potential of female mosquitoes. Overall, our findings uncover a crucial role of sugar feeding in mosquito antiviral immunity and vector competence for arboviruses. Since *Ae. aegypti* almost exclusively feed on blood in some natural settings, our findings suggest that this could increase the spread of mosquito-borne arboviral diseases.

## Introduction

Male and female adult mosquitoes regularly feed on plant nectar, thus intaking carbohydrates for their energy reserves (Foster 1995, Barredo and DeGennaro 2020). In addition, many mosquito species, including *Aedes aegypti* (*Ae. aegypti*), have evolved towards anautogeny, *i.e.* adult females require a blood meal to develop their eggs. Indeed, ingestion of blood activates neuro-endocrine and metabolic cascades leading to the synchronous development of up to hundreds of eggs which are laid two to three days after blood feeding. The blood meal is also largely used to provide proteins (*i.e.* amino acids), necessary for the synthesis of yolk proteins and for reproductive output (Clements 1992). This blood meal requirement for reproduction, and the fact that a female mosquito can take several blood meals throughout its adult life, results in *Ae. aegypti* being a vector of numerous arthropod-borne viruses (arboviruses). When a mosquito ingests a blood meal from an arbovirus-infected vertebrate host, the arbovirus initially infects the mosquito gut, then disseminates to other tissues before finally reaching the salivary glands. Once the salivary glands are infected, the arbovirus can be transmitted to a new vertebrate host during a subsequent blood meal. Arboviruses transmitted by mosquitoes, such as dengue (DENV), chikungunya (CHIKV), and Zika (ZIKV) viruses, constitute a substantial worldwide public health threat and economic burden with an increasing population at risk due to an expansion of their geographical range and an unprecedented emergence of epidemic arboviral diseases (Shepard, Undurraga et al. 2016, Gould, Pettersson et al. 2017, Wilder-Smith, Gubler et al. 2017, Weaver, Charlier et al. 2018, Messina, Brady et al. 2019).

The efficiency of mosquito-borne disease transmission under natural conditions is referred to as the vectorial capacity (Smith, Battle et al. 2012). Mathematical modelling of vectorial capacity shows that the spread of mosquito-borne pathogens is highly dependent on mosquitoes (Smith, Battle et al. 2012), including their vector competence, which is the mosquito’s ability to become infected following an infectious blood meal and subsequently transmit the pathogen (Hardy, Houk et al. 1983). Therefore, unravelling the factors which make a mosquito a competent vector will ultimately increase our fundamental understanding of mosquito-borne disease emergence and spread. In addition, it will help the development of effective and sustainable vector control strategies aimed at blocking pathogen transmission. Mosquito immunity is an important factor, among others, influencing mosquito vector competence for arboviruses (Tabachnick 2013, Bartholomay and Michel 2018, Ruckert and Ebel 2018). Mosquitoes possess different immune pathways that can control arbovirus infection (Sim, Jupatanakul et al. 2014, Cheng, Liu et al. 2016, Wu, Yu et al. 2019). This includes the NF-ĸB Toll (Xi, Ramirez et al. 2008, Anglero-Rodriguez, MacLeod et al. 2017) and immune deficiency (Imd) (Carissimo, Pondeville et al. 2015, Barletta, Nascimento-Silva et al. 2017) pathways, the Janus kinase/signal transducers and activators of transcription (JAK-STAT) pathway (Souza-Neto, Sim et al. 2009, Carissimo, Pondeville et al. 2015, Anglero-Rodriguez, MacLeod et al. 2017), and the prophenoloxidase cascade (Rodriguez-Andres, Rani et al. 2012). In addition to these immune pathways, RNA interference pathways also play an important antiviral role in mosquitoes. The exogenous small interfering (exo-siRNA) pathway in particular is a major antiviral pathway limiting the replication of many arboviruses (Olson and Blair 2015, Samuel, Adelman et al. 2018). Finally, the PIWI-interacting RNA (piRNA) pathway, and more particularly the PIWI4 protein, is involved in the regulation of arboviral replication (Schnettler, Donald et al. 2013, Varjak, Maringer et al. 2017, Tassetto, Kunitomi et al. 2019).

The particular mode of nutrition of mosquitoes linked to their extreme adaptation (*i.e.* anautogeny), alternating between sugar (carbohydrates for energy) and blood (proteins for egg development), makes them a uniquely relevant system to fundamentally understand how food intake affects the female physiology. While the influence of sugar and blood feeding on mosquito survival and reproduction, two other important determinants of vectorial capacity, have largely been investigated (Clements 1992, Foster 1995, Clements 1999, Stone and Foster 2013), the impact of sugar feeding on mosquito immunity and vector competence for arboviruses has not been analysed yet. Here, we investigated the effect of sugar feeding on immunity and susceptibility to viral infection in the main arbovirus vector, *Ae. aegypti*. We show that sugar feeding increases and maintains expression levels of genes involved in antiviral pathways in the digestive tract of females. The three major sugars naturally found in plant nectar - sucrose, glucose and fructose - can all increase antiviral gene expression. Our data show that the gut microbiota does not mediate sugar-induced immunity but partly inhibits it. Importantly, sugar intake prior to an arbovirus-infected blood meal further protects females against infection with arboviruses from different families, Semliki Forest Virus (SFV, *Alphavirus*, *Togaviridae*) and ZIKV (*Flavivirus*, *Flaviviridae*), highlighting a broad antiviral action of sugar. We further show that sugar feeding blocks arbovirus initial infection and dissemination from the gut, lowers infection prevalence and intensity, thereby decreasing the transmission potential. Overall, our findings uncover that sugar feeding is a crucial factor influencing mosquito vector competence for arboviruses, and this might affect the spread of arboviruses by *Ae. aegypti* mosquitoes.

## Results

### Sucrose feeding increases the expression levels of antiviral genes in the digestive tract of females

First, we investigated whether sugar could affect immunity in the female digestive tract. After a sugar meal, the ingested solution is initially stored in the mosquito crop. From there, the sugar solution is then periodically relocated from the crop to the midgut for digestion according to metabolic/physiological needs (Figure S1; (Foster 1995, Souza-Neto, Machado et al. 2007, Stoffolano and Haselton 2013)). After about two days (as observed in our conditions and previously described in (Foster 1995)), the crop of nearly every female is empty of all sugar solution previously ingested. Therefore, in order to compare females able to relocate and digest sugar to females that cannot, females were not given access to sucrose solution for 48 h and were then split into two groups. One group was kept without sugar (non-sugar fed, NSF). The other group was given a sucrose meal for an hour and only sugar fed (SF) females from this group were further analysed. The digestive tracts of NSF and SF females were dissected at different time points following sucrose feeding time (Figure 1A). As NSF females started to die from about 20-24 h post sucrose feeding time (*i.e.* 3 days after sugar removal), only SF females were dissected at 20, 24, and 48 h post sucrose feeding. Expression levels of representative mosquito genes involved in antiviral pathways were analysed by RT-qPCR (Figure 1B-1F). These genes include *p400* and *ago2*, involved in the siRNA pathway (Blair and Olson 2015, McFarlane, Almire et al. 2020), *piwi4* involved in the piRNA pathway (Varjak, Maringer et al. 2017, Tassetto, Kunitomi et al. 2019), *vir1*, a downstream effector of the JAK-STAT pathway (Dostert, Jouanguy et al. 2005, Souza-Neto, Sim et al. 2009), and *ppo8* involved in the phenoloxidase pathway (Zou, Shin et al. 2008, Rodriguez-Andres, Rani et al. 2012). While the gene expression levels were identical between the two populations just before the sugar feeding time (BSF, 48 h after sugar solution removed), the expression levels of *p400*, *piwi4* and *ppo8* were higher in SF females compared to NSF females at 2 h and 16 h post sucrose feeding time, with statistical significance at 16 h (Figure 1B, 1E, 1F). The expression levels of *ago2* were unchanged (Figure 1C) and the expression levels of *vir1*, although slightly higher in SF females at 2 h post sucrose meal, were lower in SF females at 16 h (Figure 1D). The expression of all analysed genes was upregulated in SF females at 24 and 48 h after sugar feeding compared to before, with *p400*, *piwi4* and *ppo8* peaking at 24 h (Figure 1B, 1E, 1F), and *ago2* and *vir1* at 48 h (Figure 1C, 1D). Concomitantly with an increase of immune gene expression levels in SF females over time (significant for all genes except *ago2*), a decrease of immune expression levels occurred in NSF females (significant for *p400* and *piwi4*). The expression levels of most genes in SF females were oscillating from sugar feeding to 48 h, with an alternance of increase and decrease of levels compared to the previous time point (Figure 1). This is not linked to circadian rhythm as digestive tracts sampled either before sugar feeding time or at 24 and 48 h after sugar feeding were dissected at the same time of day. Instead, this is more consistent with sugar relocation from the crop to the gut happening periodically (Foster 1995, Stoffolano and Haselton 2013). The activation of immune gene expression was well due to sucrose, as the expression levels of *p400*, *piwi4* and *ppo8* in females fed with 2% sucrose were intermediary between the ones of NSF females and females fed with 10% sucrose (Figure S2). Altogether, these results show that sucrose increases and maintains antiviral gene expression levels in the gut of *Ae. aegypti* females.

**Figure 1.**
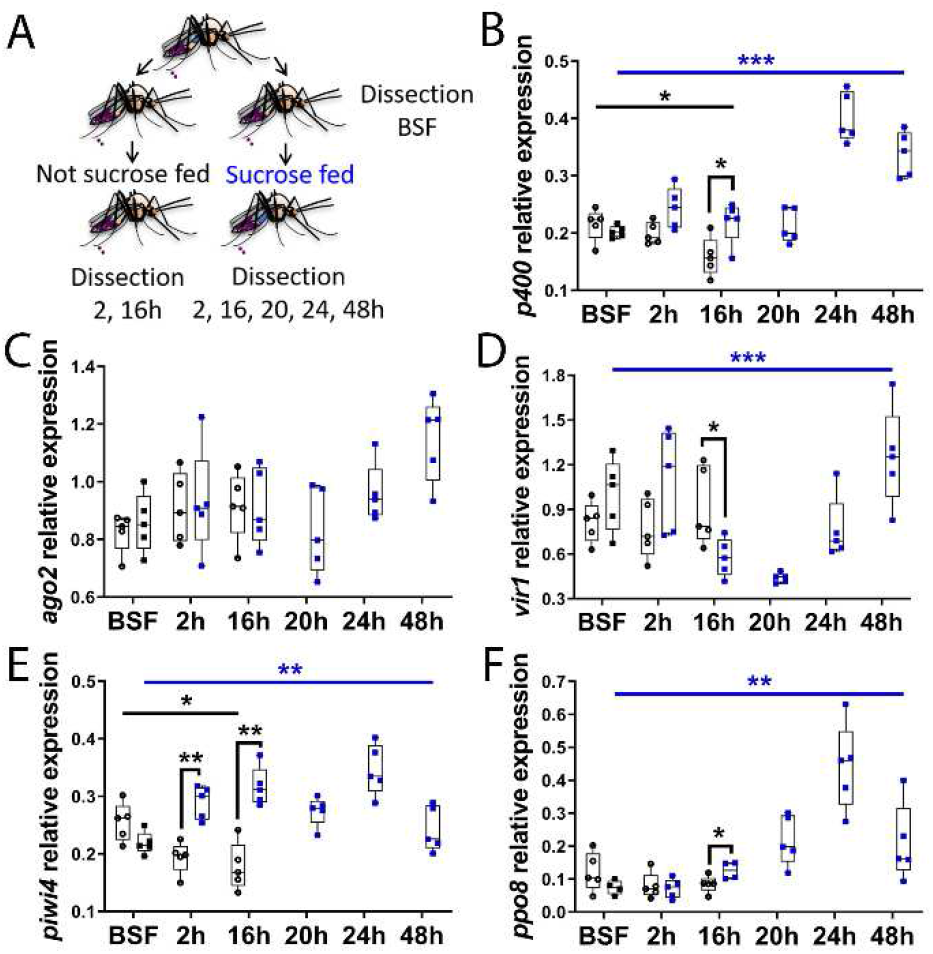
Sucrose increases the expression levels of antiviral genes in the digestive tract of the female *Ae. aegypti*. (A) Schematic of experimental design. Females that did not have access to sucrose for 48 h were fed or not with 10% sucrose solution. Digestive tracts from both populations were dissected just before sucrose feeding (BSF) and at 2 and 16 h post sucrose feeding time, and at 20, 24 and 48 h for the sucrose fed females. (B to F) RNA transcript levels of (B) *p400*, (C) *ago2*, (D) *vir1*, (E) *piwi4* and (F) *ppo8* (black empty dots and black squares, non-sucrose fed; blue squares, sucrose fed). Box plots display the minimum, first quartile, median, third quartile, and maximum relative expression levels. Data were analysed by Mann-Whitney test (non-sucrose vs sucrose fed at each time point, BSF, 2 and 16 h) and by Kruskal-Wallis test (all times, for non-sucrose fed and sucrose fed, black and blue bars respectively showing p value summary). N = 5 pools of 5 digestive tracts per condition. Only p values < 0.05 are shown. *, p value < 0.05; ***, p value <0.001.

### Glucose and fructose increase the expression of antiviral genes in the digestive tract of females

The disaccharide sucrose ingested by mosquitoes is hydrolysed into the monosaccharides glucose and fructose by α-glucosidases secreted in the saliva and bound to midgut epithelium membranes (Marinotti and James 1990, Burkett, Carlson et al. 1998, Souza-Neto, Machado et al. 2007). These monosaccharides, along with sucrose, are the most abundant sugars in nectar and are present in the natural sugar meals ingested by mosquitoes (Burkett, Carlson et al. 1998). Therefore, we asked whether glucose and/or fructose were responsible for the sucrose-mediated increase of antiviral gene expression. Hence, females were not fed, or fed with either sucrose, glucose or fructose. Digestive tracts were dissected at 16 h post sugar feeding (Figure 2A), a time point at which the sucrose-induced antiviral gene expression was observable for *p400*, *piwi4* and *ppo8* in the previous time course experiment (Figure 1). Expression levels of these antiviral genes in the digestive tract of the females were analysed by RT-qPCR (Figure 2B-2D). As with sucrose, the expression levels of *p400*, *piwi4* and *ppo8* were significantly upregulated in the gut of the females fed with glucose or fructose, compared to NSF females. Thus, sucrose, glucose and fructose can increase and maintain the expression of antiviral genes in the digestive tract of *Ae. aegypti* females.

**Figure 2.**
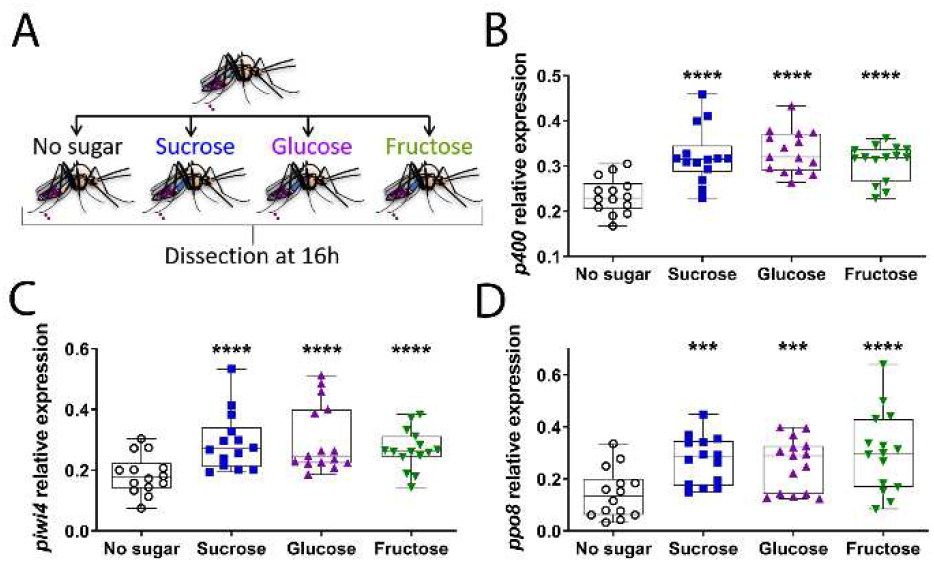
Sucrose, glucose and fructose increase the expression levels of antiviral genes in the digestive tract of the female *Ae. aegypti*. (A) Schematic of experimental design. Females that did not have access to sucrose for 48 h were either not sugar fed or fed with 10% sucrose, 10% glucose or 10% fructose solution. Digestive tracts were dissected 16 h post sugar feeding time. (B to D) RNA transcript levels of (B) *p400*, (C) *piwi4* and (D) *ppo8*. Box plots display the minimum, first quartile, median, third quartile, and maximum relative expression levels. Data from three separate experiments were combined after verifying the lack of a detectable experiment effect. N = 15 pools of 5 digestive tracts per condition. Statistical significance of the sugar feeding effect was assessed with an analysis of variance followed by a Fisher’s multiple comparison test (Sugar vs No sugar). ***, p value < 0.001; ****, p value <0.0001.

### Sugar-induced antiviral gene expression is not mediated by but inhibited by gut bacteria

Diet, including sugar, can induce gut microbial shifts (both in terms of diversity and abundance) in different hosts (Singh, Chang et al. 2017, Harris, de Roode et al. 2019, Satokari 2020). Since endogenous gut bacteria can modulate immune gene expression in mosquitoes (Jupatanakul, Sim et al. 2014, Rodgers, Gendrin et al. 2017), we hypothesised that the increase of immune gene expression levels in the digestive tract following sugar feeding might be influenced by gut bacteria. To investigate this, the effect of sugar feeding on immune gene expression levels was compared between control mosquitoes and mosquitoes from which the gut bacteria had been eliminated through antibiotic treatment (Figure 3A, Figure S3). As the gut microbiota can activate genes regulated by the Toll and Imd immune pathways, including antimicrobial peptides (AMPs) (Xi, Ramirez et al. 2008, Ramirez, Souza-Neto et al. 2012, Barletta, Nascimento-Silva et al. 2017), the expression of two AMPs, *cecropin* (*cecD*) and *defensin* (*defE*), was analysed in addition to previously assessed antiviral genes at 16 h post sucrose feeding time (Figure 3B-H). As previously observed, sucrose feeding significantly increased the expression levels of immune genes (*e.g. p400*, *piwi4* and *ppo8*) in the gut of control females (Figure 3B, 3E, 3F). Sugar feeding also significantly increased the expression of *cecD* (Figure 3G). Elimination of the gut bacteria in NSF females had little to no effect on immune gene expression except for *cecD* (Figure 3G), which was as expected expressed at lower levels in aseptic mosquitoes. Intriguingly, antibiotic treatment did not abolish the sugar-induced increase of immune gene expression. Sugar feeding drastically increased the expression of *p400*, *piwi4* and *ppo8* in the gut of aseptic females. Most of the genes were expressed at higher levels in the gut of sugar fed aseptic females compared to sugar fed control females, except *cecD* whose expression levels again decreased upon antibiotic treatment. Analyses revealed a significant overall effect of both sugar and/or antibiotic treatments for most of the genes as well as a significant interaction between sugar and antibiotic treatments for *p400* and *piwi4* (two-way ANOVA statistical significance of treatments, Figure S4). Although not significant, the expression levels of the two AMPs, *cecD* and *defE*, were also increased upon sugar feeding in the gut of aseptic females. Altogether, our results show that gut bacteria do not mediate the increase of immunity after a sugar meal, but partially inhibit the sugar-induced immune gene activation in the gut of the female *Ae. aegypti* mosquito.

**Figure 3.**
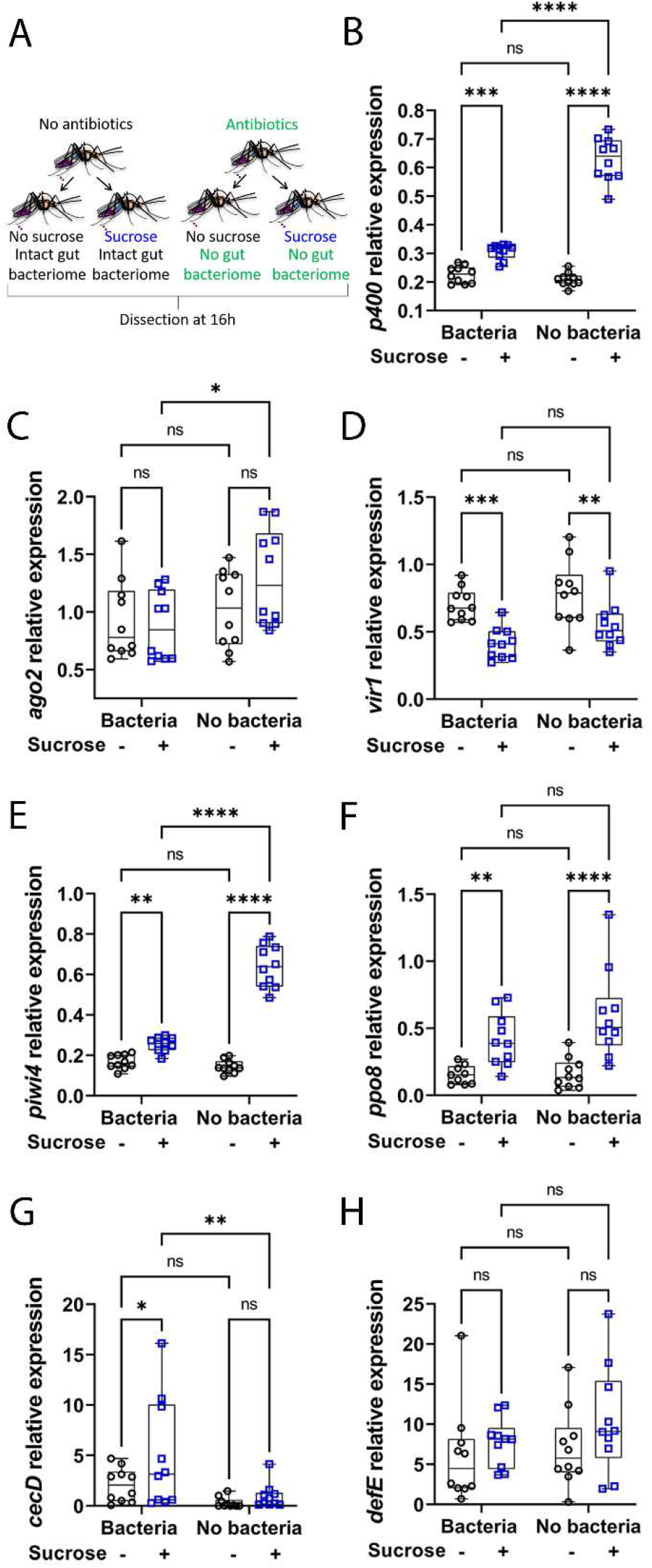
Microbiota partly inhibits sugar-induced immunity in the digestive tract of the female *Ae. aegypti*. (A) Schematic of experimental design. Females, previously treated or not with antibiotics, did not have access to sucrose for 48 h and were either not fed or fed with 10% sucrose. Digestive tracts were dissected 16 h post sugar feeding time. (B to H) RNA transcript levels of (B) *p400*, (C) *ago2*, (D) *vir1*, (E) *piwi4*, (F) *ppo8*, (G) *cecD* and (H) *defE*. Box plots display the minimum, first quartile, median, third quartile, and maximum relative expression levels. Data from two separate experiments were combined after verifying the lack of a detectable experiment effect. N = 10 pools of 5 digestive tracts per condition. Statistical significance of the treatments effect was assessed with a two-way ANOVA (statistical analysis summary of treatment effect and interaction between treatments given in Figure S4). Pair-wise comparisons shown on the graphs were obtained with a post hoc Fisher’s multiple comparison test (No sucrose vs Sucrose, Bacteria vs No bacteria). ns, p value > 0.05; *, p value < 0.05; **, p value < 0.01; ***, p value <0.001; ****, p value <0.0001.

### Sucrose intake prior to an arbovirus-infected blood meal protects females against alphavirus infection and dissemination from the gut

The gut is the first barrier to be overcome by an arbovirus and is considered to be a major determinant of arbovirus infection and mosquito vector competence (Franz, Kantor et al. 2015, Cheng, Liu et al. 2016). After an infectious blood meal, arboviruses must rapidly infect the midgut cells before the formation of the peritrophic matrix and to avoid the proteasic environment inherent to blood digestion (Sanders, Evans et al. 2003, Franz, Kantor et al. 2015). We showed that the immune gene expression in the digestive tract is enhanced following sucrose feeding and even more in the absence of the gut bacterial flora (Figure 3). Since the genes analysed here are involved in antiviral pathways in *Ae. aegypti*, we therefore reasoned that a sugar meal before an infectious blood meal may decrease initial arbovirus midgut infection and replication and further dissemination from the gut. To investigate this, NSF or SF females were given a blood meal infected with the arbovirus Semliki Forest virus (SFV), a prototype alphavirus of the *Togaviridae* family, 16 h after the sugar feeding time. As sugar-induced immunity is stronger in aseptic females, the effect of sugar feeding on infection outcome was analysed in both control and aseptic females to determine whether effects would be correlated to the strength of sugar-induced immune activation (Figure 4A). Analysis of immune gene expression in the digestive tracts at 6 h after the blood meal (Figure 4B-H), revealed an overall significant effect of the sugar treatment on *p400*, *ago2*, *piwi4* and *ppo8* and a significant interaction between sugar and antibiotic treatment for *p400*, *ago2*, *vir1* and *piwi4* (two-way ANOVA statistical significance of treatments, Figure S4). Although 6 h after blood meal, sugar-mediated induction of immune gene expression was no longer detectable in the gut of females with intact bacterial flora, it was still observed in the gut of antibiotic-treated females (Figure 4B-H). Since blood meal provokes a reduction in sugar response in the gut (Foster 1995, Sanders, Evans et al. 2003), it is possible that gene expression levels in SF females at the time of blood feeding (16 h post sugar feeding, Figure 3) progressively decrease after a blood meal. As the effect of sugar is stronger at the time of blood feeding (16 h post sugar feeding, Figure 3) in the gut of aseptic females compared to control ones, this may be delayed in aseptic females.

**Figure 4.**
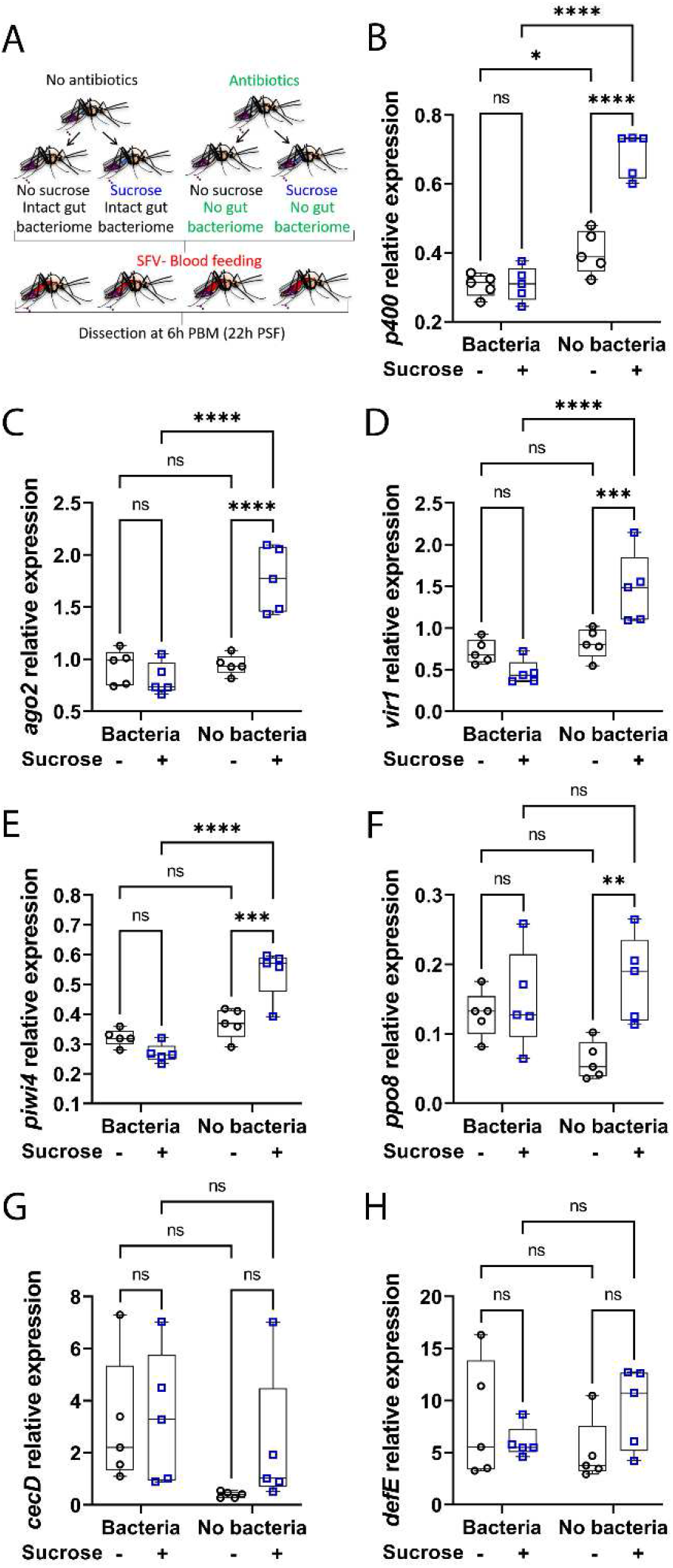
Sugar feeding prior to a blood meal increases antiviral gene expression in blood fed females. (A) Schematic of experimental design. Females, previously treated or not with antibiotics, did not have access to sucrose for 48 h and were either not fed or fed with 10% sucrose. Females received an SFV-infected blood meal at a titre of 7.8 × 10^7^ PFU/mL 16 h post sugar feeding time and digestive tracts were dissected 6 h after the blood meal. PBM, post blood meal; PSF, post sugar feeding. (B to H) RNA transcript levels of (B) *p400*, (C) *ago2*, (D) *vir1*, (E) *piwi4*, (F) *ppo8*, (G) *cecD* and (H) *defE*. Box plots display the minimum, first quartile, median, third quartile, and maximum relative expression levels. N = 5 pools of 5 digestive tracts per condition. Statistical significance of the treatments effect was assessed with a two-way ANOVA (statistical analysis summary of treatment effect and interaction between treatments given in Figure S4). Pair-wise comparisons shown on the graphs were obtained with a post hoc Fisher’s multiple comparison test (No sucrose vs Sucrose, Bacteria vs No bacteria). ns, p value > 0.05; *, p value < 0.05; **, p value < 0.01; ***, p value <0.001; ****, p value <0.0001.

Importantly, although the quantity of initial infectious particles in females was similar between the different groups (Figure 5A), sugar feeding prior to the infectious blood meal significantly decreased SFV infection (gut), dissemination (body) and transmission potential (head, as a proxy of virus in salivary glands) outcome at 4 days post infection (Figure 5B-E, Figure S4). In septic females, sugar feeding reduced SFV titres in infected guts, bodies and heads (Figure 5B-D). While antibiotic treatment had little to no effect on infection titres and prevalence in NSF females, antibiotic treatment led to an even more pronounced effect of sugar on infection outcome (Figure 5B-E). Although there was only a trend for a reduction in SFV titres in infected body parts from aseptic females (Figure 5B-D), sugar feeding strongly reduced prevalence of gut infection and dissemination to the body and head (Figure 5E). Thus, when not partially inhibited by the gut bacteria, sugar feeding increased resistance to infection, with only a small proportion of engorged females still showing a transmission potential (20%), *e.g.* virus dissemination to the head (Figure 5E). Infection prevalence analysis shows that once virus was in the body, sugar treatment no longer had an effect on dissemination (Figure 5E). Therefore, the impact of sugar feeding on transmission potential prevalence was only due to an effect of sugar on initial gut infection and dissemination from the gut, demonstrating that the gut is a crucial determinant in mosquito vector competence.

**Figure 5.**
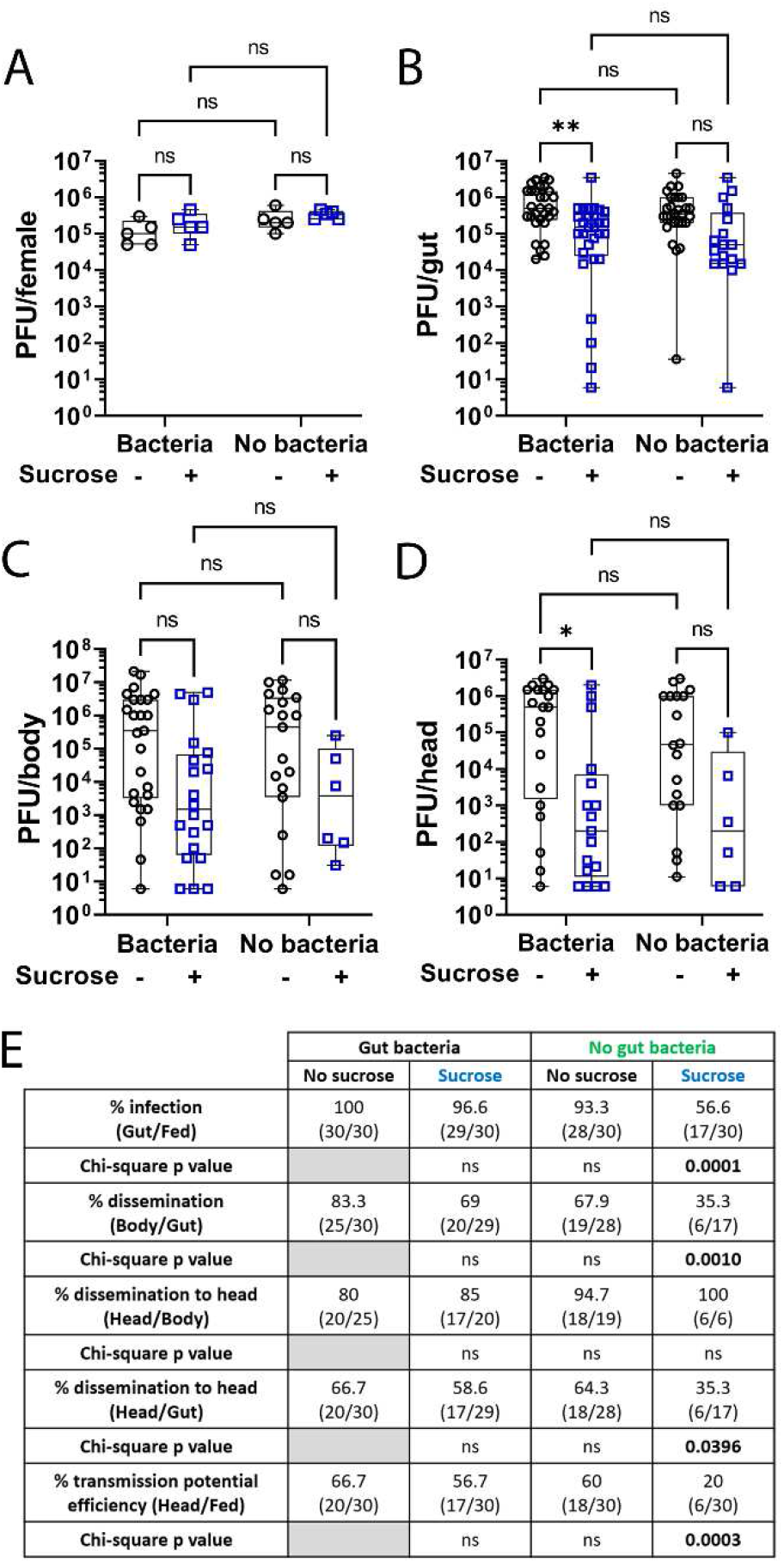
Sugar feeding protects the female mosquito *Ae. aegypti* against SFV infection. Females were treated as in Figure 4A. All females were given access to sucrose solution 30 h after the infectious blood meal. (A) Virus titration by plaque assays of individual females sacrificed straight after blood meal to assess infectious particles ingested. N = 5 per condition. (B to D) Virus titration by plaque assays of individual (B) guts, (C) bodies and (D) heads sampled four days post infection. N = 30 tissues per condition. Only infected samples (with less than 5 PFU per tissue, *i.e.* no plaque in two replicate wells at the first dilution) were plotted on the graphs. Box plots display the minimum, first quartile, median, third quartile, and maximum. Data were analysed using two-way ANOVA (statistical analysis summary of treatment effect and interaction between treatments given in Figure S4). Pair-wise comparisons shown on the graphs were obtained with a post hoc Fisher’s multiple comparison test (No sucrose vs Sucrose, Bacteria vs No bacteria). (E) SFV infection, dissemination and transmission potential prevalence (in percentage and numbers in brackets). The p values indicate statistical significance of the treatments effect on prevalence assessed with a Chi-square test (compared to No bacteria - No sucrose group). ns, p value > 0.05.

### Sucrose intake protects females against flavivirus infection

To determine if sugar feeding and its interaction with the gut bacteria could impact on other arboviruses, the experiment was repeated with ZIKV (*Flavivirus*, *Flaviviridae*). Here, whole females were sampled at 6 days post infection and viral RNA levels determined by RT-qPCR (Figure 6). Sugar and antibiotic treatments as well as their interaction had little effect on ZIKV RNA levels in infected females (Figure 6A, Figure S4). Indeed, viral RNA levels were only slightly reduced upon sugar feeding in both control and aseptic infected females (Figure 6A). Nonetheless, ZIKV infection prevalence was dramatically reduced upon sugar feeding in both control and aseptic females, and more strongly in the latter ones (Figure 6B). Interestingly, antibiotic treatment also decreased ZIKV infection prevalence in NSF females (Figure 6B), suggesting that the gut microbiota could also limit ZIKV infection independently of the sugar effect. Flaviviruses take longer to disseminate from the gut (Ryckebusch, Berthet et al. 2017, Winokur, Main et al. 2020) than alphaviruses (Dubrulle, Mousson et al. 2009). Since females from all groups had access to sugar 30 h after the blood meal, and that sugar-induced immunity is stronger in aseptic females, it is possible that less of these females were infected thanks to an additional protective role of sugar imbibed after the blood meal. Altogether, our findings reveal that sugar feeding before an infectious blood meal protects *Ae. aegypti* females against different arboviruses.

**Figure 6.**
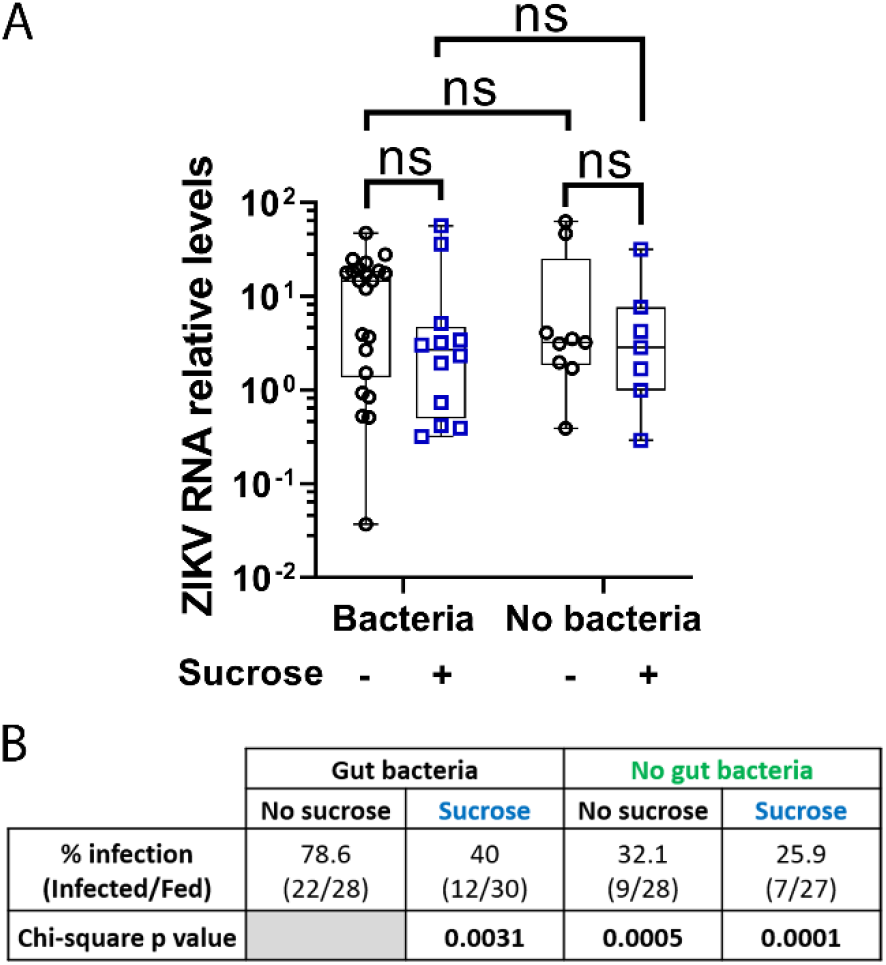
Sugar feeding protects the female mosquito *Ae. aegypti* against ZIKV infection. Females were treated as in Figure 4A except that females received a ZIKV-infected blood meal at a titre of 5.3 × 10^6^ PFU/mL. All females were given access to sucrose solution 30 h after infectious blood meal. (A) ZIKV RNA levels, relative to *S7 ribosomal protein* transcript, in individual whole females assessed by RT-qPCR. Data were analysed as described (Taylor, Nadeau et al. 2019) so as the RQ geomean of the group ‘No antibiotics-No sucrose’ is 1. N = 27 to 30 females per condition. Only ‘infected’ samples (with Ct less than 30) were plotted on the graphs. Box plots display the minimum, first quartile, median, third quartile, and maximum RQ values. Log2-transformed values of RQ values were analysed using two-way ANOVA (statistical analysis summary of treatment effect and interaction between treatments given in Figure S3). Pair-wise comparisons shown on the graphs were obtained with a post hoc Fisher’s multiple comparison test (No sucrose vs Sucrose, Bacteria vs No bacteria). (B) ZIKV infection prevalence (in percentage and numbers in brackets). The p values indicate statistical significance of the treatment effect on prevalence assessed with a Chi-square test (compared to No bacteria-No sucrose group).

## Discussion

Diet profoundly influences all aspect of animal biology and physiology. In particular nutrition is a key regulator of health and disease outcomes through interactions with the immune system (Nobs, Zmora et al. 2020). For instance, nutrition can impact arbovirus virulence in mice (Weger-Lucarelli, Carrau et al. 2019). So far, the effect of diet on immunity and health has been mostly studied in humans and model organisms such as mice and *Drosophila melanogaster*. However, assessing the impact of diet in biomedically relevant species, such as insect disease vectors, could inform on the role of nutrition in the spread of diseases and support the design of control strategies. Here, our findings unveil that sugar feeding increases and maintains antiviral immunity in the digestive tract, and further protects the arbovirus vector *Ae. aegypti* against arbovirus infection.

Consistent with our findings that in *Ae. aegypti* females, sugar feeding promotes the expression of immune genes, including *ppo8* which is involved in the melanisation cascade, *Anopheles* and *Culex* females are dependent on a sugar meal to mount an effective melanisation response (Koella and Sorense 2002, Ferguson, Beckett et al. 2019). A recent study has also found a positive effect of carbohydrates on immune gene expression in *Drosophila* (Ponton, Morimoto et al. 2020). While studies investigating the role of diet carbohydrates on insect immunity are very scarce, some reports have highlighted a negative effect of protein and/or a positive effect of carbohydrates on resistance to infection. For instance, feeding on a high-carbohydrate diet increases survival following infection with a gut pathogen in *Drosophila* (Galenza, Hutchinson et al. 2016) and fungal infection in ants (Kay, Bruning et al. 2014). Moreover, inhibition of glucose utilisation is lethal to mice under influenza infection (Wang, Huen et al. 2016). Although studies in more models are warranted, the protective role of sugar against infection may be conserved across evolution.

The genes analysed here are known to be antiviral or to be involved in antiviral immune pathways in mosquitoes including *Ae. aegypti* (Sim, Jupatanakul et al. 2014, Cheng, Liu et al. 2016, Wu, Yu et al. 2019). Besides, it has been previously shown that the basal levels of immune-related transcripts in the mosquito gut influence susceptibility to infection and vector competence for arboviruses (Sim, Jupatanakul et al. 2013). Moreover, given that i) sugar feeding leads to inversely correlated changes in immune gene expression (increase) and virus titres/prevalence (decrease) and ii) elimination of the gut bacteria results in a higher sugar-induced immune gene activation and a stronger protection against arbovirus infection, it is likely that the effect of a sugar meal prior to an infectious blood meal on viral infection is mediated, at least in part, by sugar-induced antiviral immunity. Notably, the decrease of transmission potential efficiency in females previously fed with sugar (proportion of heads infected on blood fed females) was only due to a reduction in initial gut infection and further dissemination from the gut. Therefore, the early stage of infection in the gut and initial susceptibility to arboviruses is a crucial determinant of mosquito vector competence.

Although it is widely reported that in nature, male and female mosquitoes feed on sugar sources to maintain their energy reserves, the degree to which *Aedes* female mosquitoes require sugar meals remains unclear. Some studies have shown that *Aedes* mosquitoes seek nectar throughout their lifetime (Andersson 1990, Smith and Gadawski 1994, Martinez-Ibarra, Rodriguez et al. 1997), while others have found that *Aedes* females feed rarely on sugar (Edman, Strickman et al. 1992) and almost exclusively feed on blood (Scott, Chow et al. 1993). Because non-sylvatic *Ae. aegypti* mosquitoes are adapted to human environment and often found indoors, they appear to bite more frequently and throughout a single gonotrophic cycle, acquiring multiple blood meals that are used for reproduction, energy, and survival (Scott, Chow et al. 1993, Scott, Amerasinghe et al. 2000, Reyes-Villanueva 2004, Farjana and Tuno 2013). Since *Ae. aegypti* female mosquitoes fed on blood alone survive as long as females fed on sugar alone (Canyon, Hii et al. 1999) or blood with access to sugar (Costero, Edman et al. 1998), it suggests that sugar may be dispensable. Whether mosquito females take sugar meals or not appears to be dependent on different factors, including the availability of sugar sources (Ignell, Okawa et al. 2010). Overall, the absence of sugar feeding in some natural settings may explain, in part, the high susceptibility and vector competence of *Ae. aegypti*.

Sugar feeding can also influence mosquito blood feeding frequency, survival and reproduction, which can impact on vectorial capacity. However, whether sugar feeding affects the vectorial capacity and pathogen transmission rate is still unclear (Foster 1995, Stone and Foster 2013, Barredo and DeGennaro 2020). In *Anopheles* mosquitoes, it has been estimated that vectorial capacity could be increased in nectar-rich environments because of an increased survival and reproductive rate (Ebrahimi, Jackson et al. 2018). However, the precise role of sugar feeding in vector competence would need to be investigated in these mosquito species to account for this parameter in the estimation of the impact of sugar on vectorial capacity. In *Ae. aegypti*, sugar feeding may not increase reproduction and survival, as feeding only on human blood increases these two life traits compared to feeding on blood and sugar (Costero, Edman et al. 1998, Harrington, Edman et al. 2001). In addition, sugar fed *Ae. aegypti* females have a reduced avidity for a blood meal (Jones and Madhukar 1976). Taken together, this suggests that sugar feeding, by decreasing vector competence, could reduce vectorial capacity in *Ae. aegypti*. In the urge of implementing new strategies for mosquito vector control, the use of sugar baits, supplemented with insecticides, have been developed and have proven to be effective at attracting female mosquitoes outdoor as well as indoor, and regardless of the availability of natural sugar sources (Beier, Müller et al. 2012, Qualls, Müller et al. 2015, Fiorenzano, Koehler et al. 2017, Sissoko, Junnila et al. 2019). Although more experimental and modelling studies of the influence of sugar on the vectorial capacity of arbovirus vectors are necessary, our findings highlight that basic sugar baits could constitute an environmentally friendly, sustainable, and cost-effective method to control arbovirus transmission.

In *Drosophila*, sugar-mediated survival to intestinal infection is independent of the bacterial flora (Galenza, Hutchinson et al. 2016). Similarly, we found that sugar-induced antiviral immunity and protection against arbovirus infection were not mediated by the gut bacteria. Sugar-mediated effects were stronger in aseptic mosquitoes, showing that the beneficial action of sugar on antiviral immunity and protection against arbovirus infection can in part be inhibited by gut bacteria. Studies have highlighted the ability of mosquito-associated bacteria to reduce (Xi, Ramirez et al. 2008, Ramirez, Souza-Neto et al. 2012, Barletta, Nascimento-Silva et al. 2017) or, similarly to our observations, increase (Apte-Deshpande, Paingankar et al. 2012, Carissimo, Pondeville et al. 2015, Wu, Sun et al. 2019) vector competence for arboviruses. Different mechanisms, such as the modulation of immunity, have been proposed to explain the influence of bacteria on mosquito vector competence (Jupatanakul, Sim et al. 2014). In our study, gut bacteria did not affect immunity in NSF females but only in SF females. Although it cannot be excluded that bacteria inhibition is independent of the sugar activation effect, *i.e.* if basal levels of immunity in NSF females were too low to observe inhibition by bacteria, it suggests that microbiota could directly inhibit the sugar induction of immune gene expression in SF females.

Gaining a better understanding of how nutrition influences immunity and resistance to infection in biomedically relevant species is a significant challenge. Overall, our study unveils the nutritional regulations of innate immune gene expression and resistance to arbovirus infection in mosquitoes. Analysis of vector competence is often done in laboratories, where females can feed *ad libitum* on a sugar solution. It is interesting to note that it is a common practice to remove the sugar solution a few hours to a day before a blood meal (infectious or not). Therefore, our results also shed light on unsuspected factors that could influence and introduce variations in the outcome of laboratory-based infection experiments. More importantly, our findings increase our fundamental understanding of the factors influencing vector competence. The role of sugar in vector competence might affect transmission rates of mosquito-borne arboviral diseases and therefore should be included in epidemiological models. Ultimately, this should ultimately help the development and application of vector control strategies, such as sugar baits, aimed at reducing arbovirus transmission.

## Material and Methods

### Mosquito rearing

*Ae. aegypti* Paea strain (a gift of Dr A-B. Failloux, Institut Pasteur, France) was reared at 28°C and 80% humidity conditions with a 12 h light/dark cycle. Larvae were reared in water and fed with dry cat food (Friskies). Emerging adult mosquitoes were maintained on a 10% (w/vol) sucrose solution *ad libitum*. Females were fed with heparinised rabbit blood (Envigo, UK) for 1 h using a 37°C Hemotek system (Hemotek Ltd, Blackburn, UK).

### Sugar, antibiotic and blood feeding treatments

From emergence, adult mosquitoes were fed with unlimited access to 10% sucrose solution supplemented or not with 200 U/mL Penicillin/Streptomycin (Gibco) and 200 μg/mL Gentamicin (Invitrogen). Sucrose solution containing antibiotics was replaced every day. Sucrose solution was removed from the cages at 7 days post emergence for 48 h before females were given, or not, a 10% sucrose meal (Fisher Chemical) supplemented or not with antibiotics (same concentration as above), a 2% sucrose meal (Fisher Chemical), a 10% glucose (Sigma-Aldrich) solution or a 10% fructose (Sigma-Aldrich) solution, all containing 1 mg/ml of non-absorbable Erioglaucine disodium salt (Sigma-Aldrich) for 1 h using a 37°C Hemotek system (Hemotek Ltd, Blackburn, UK). Sugar fed females were isolated after feeding and further kept. To analyse immune gene expression in BF females, females were blood fed 16 h post sucrose feeding time with heparinised rabbit blood (ENVIGO) for 1 h using a 37°C Hemotek system (Hemotek Ltd, Blackburn, UK). To analyse immune gene expression, the digestive tracts of female mosquitoes (N = 5 pools of 5 tissues per condition and per independent experiment) were dissected in PBS 1X with 0.05% Tween 20, sampled in RNA later (Sigma-Aldrich) and stored at 4°C until RNA extraction.

### Antibiotic efficacy validation

Antibiotic efficacy was controlled by both Luria-Bertani (LB) plating 16 h post sucrose feeding time and 16S qPCR 18 h post sucrose feeding time. Prior to dissection of the digestive tracts used to control the efficacy of antibiotics, females were washed for 10 sec with three consecutive baths of 70% ethanol and one bath of PBS 1X. For LB plating, five digestive tracts were pooled (per condition and per independent experiment) in 200 μL of PBS 1X and homogenized (Precellys 24, Bertin Technologies) at 6500g for 30 sec. Colony forming units were determined by plating the homogenate of the digestive tracts with series dilution on LB-agar medium under aseptic conditions and incubated at 37°C for 24 h. For 16S qPCR, five digestive tracts (per condition and per independent experiment) were pooled in 180 μL PBS 1X and were homogenized at 6500g for 30 sec. Extraction of gDNA was performed using the DNeasy blood and tissue kit (Qiagen) following the manufacturer protocol. gDNA concentration as well as optic densities (OD) 260/280 and 260/230 was measured using a nanodrop and gDNA was aliquoted and stored at −20°C until qPCR.

### Cell culture

BHK-21 cells (a kind gift of Prof. R. M. Elliott, MRC-University of Glasgow Centre for Virus Research, UK) were cultured in GMEM (Gibco), 10% FBS, 10% TPB (Gibco) and 83 U/ml penicillin/streptomycin (Gibco). A549 NPro cells (a kind gift of Prof. R. E. Randall, University of St. Andrews, UK) were maintained in DMEM (Gibco) with 10% FBS, and expression of bovine viral diarrhea virus-derived NPro, which targets IRF-3 for degradation (Hilton, Moganeradj et al. 2006, Hale, Knebel et al. 2009) was maintained by the addition of 2 μg/ml puromycin (Gibco). Vero E6 cells (ATCC#CRL-1586; a kind gift of Prof. M. Bouloy, Institut Pasteur, France) were maintained in DMEM (Gibco) supplemented with 10% FBS. All cells were maintained at 37°C in a humidified incubator with 5% CO_2_.

### Virus production

SFV4 was rescued from plasmid pCMV-SFV4 as described previously (Ulper, Sarand et al. 2008). Briefly, the plasmid was transfected using Lipofectamine 2000 (Thermo Fisher Scientific) into BHK-21 cells grown in GMEM with 2% FBS and 10% TPB at 37°C with 5% CO_2_. Harvested virus stock was amplified in BHK-21 cells in GMEM supplemented with 2% FBS and 10% TPB for 2 days; at harvest, the cellular debris was spun down by centrifugation at 4000 rpm for 10 min, the obtained virus stock was topped up with FBS to 10% and sodium bicarbonate added (1 ml of 7.5% solution per 20 ml of stock). The virus stock was aliquoted and stored at −80°C and used in subsequent experiments. ZIKV virus, rescued from pCCL-SP6-ZIKVwt (Mutso, Saul et al. 2017), was obtained from Jamie Royle (University of Glasgow Centre for Virus Research). The stock was amplified in A549 NPro cells for 8 days in DMEM supplemented with 2% FBS, and 25 mM HEPES. Upon harvest, the supernatant was clarified by centrifugation at 4000 rpm for 10 min. The supernatant was then concentrated on Amicon Ultra 15 filter columns (Merck Millipore). The obtained virus stock was topped up with FBS to 10% and sodium bicarbonate added. The virus stock was aliquoted and stored at −80°C.

### SFV and ZIKV infections

Females treated with sucrose supplemented (or not) with antibiotics since emergence were not given access to sucrose solution for about 48 h before receiving (or not) a 10% sucrose solution supplemented (or not) with antibiotics. An SFV- or ZIKV-infected blood meal was offered to females at 16 h post sucrose feeding time. To prepare the infectious blood meal, fresh rabbit blood (Envigo) was washed with PBS to remove white blood cells and serum. PBS was then added to the red blood cell fraction to return to the initial blood volume. The infectious blood meal was prepared with 2/3 of washed blood and 1/3 of virus-containing culture medium to give a final titre of 7.8 × 10^7^ PFU/mL for SFV or 5.3 × 10^6^ PFU/mL for ZIKV, supplemented with 2 mM ATP. Only engorged females were further kept at 28°C and 80% humidity. Access to 10% sucrose solution *ad libitum* was permitted from 30 h post infectious blood meal. SFV-infected females (N = 5 individual females per condition and per independent experiment) were sampled just after the infectious blood meal, homogenized in 100 μL of GMEM (Gibco) containing 10% FBS and stored at −80°C until titration to assess ingested virus quantity. To analyse immune gene expression, the digestive tracts of SFV-infected female mosquitoes (N = 5 pools of 5 digestive tracts per condition and per independent experiment) were dissected in PBS with 0.05% Tween 20, sampled in RNA later (Sigma-Aldrich) and stored at 4°C until RNA extraction. SFV-infected females were sampled four days post infection (N = 30 per condition and per independent experiment). Heads, digestive tracts and the remains of bodies were individually homogenized in 100 μL of GMEM (Invitrogen) containing 10% FBS and stored at −80°C until titration. Whole ZIKV-infected females were sampled six days post infection, homogenized individually (N = 30 per condition and per independent experiment) in 500 μl of TRIzol (Invitrogen) and stored at −80°C until RNA extraction.

### Virus titration

SFV4 stock was titrated by plaque assay on BHK-21 cells. For titration, 10-fold serial dilutions of virus stock were prepared in GMEM containing 2% FBS. Following an hour-long incubation with inoculum, the cells were overlaid with 1X MEM (Gibco) containing 2% FBS and 0.6% Avicel (FMC Biopolymer). The cells were incubated for 2 days at 37°C with 5% CO_2_, and thereafter fixed using equal volume of 8% formaldehyde solution for 1 h. Plaques were stained using 0.2% toluidine blue (Sigma-Aldrich). The titre of infectious particles is expressed as plaque forming units per ml (PFU/mL). ZIKV stock was titrated by plaque assay on Vero E6 cells. For titration, 10-fold serial dilutions of virus stock were prepared in PBS containing 2% FBS. Following an hour-long incubation with inoculum, cells were overlaid with 1X MEM (Gibco) containing 2% FBS and 0.6% Avicel. Cells were incubated for 5 days at 37°C with 5% CO_2_, and thereafter fixed using equal volume of 8% formaldehyde solution for 1 h. Plaques were stained using crystal violet solution (20% (v/v) ethanol, 1% (v/v) methanol, 0.1% (w/v) methyl violet (Fisher Scientific), diluted in H2O). Titre of infectious particles is expressed as PFU/mL. Mosquito samples infected with SFV were serially diluted in GMEM containing 2% FBS and 1X antibiotic-antimycotic solution (Final concentration: 100 U/mL penicillin, 0.1 g/mL streptomycin, 250 ng/mL amphotericin B, Sigma-Aldrich) and inoculated onto BHK-21 cells seeded the day before at 1.5 × 10^5^ cells/ml on 24-well plates (two replicate wells per sample). Plates were incubated for 1 h at 37°C with 5% CO_2_ and were then overlaid with 1X MEM (Gibco) containing 2% FBS and 0.6% Avicel for 2 days. Plates were fixed with an equal volume of 10% Formalin (Sigma-Aldrich) and plaques were revealed using 0.2% toluidine blue (Sigma-Aldrich). The average number of plaques (from 2 wells) was multiplied by the dilution factor to obtain the titre of infectious particles as PFU/tissue. Samples were considered negative when no plaque was obtained in the two replicate wells (*i.e* less than 5 PFU/sample).

### RNA extraction and reverse-transcription

RNA later was removed from the samples. Samples were homogenized (Precellys 24, Bertin Technologies) in 1 mL of TRIzol (Invitrogen), or in 600 μL of buffer RLT (Qiagen) for digestive tracts containing blood, at 6500g for 30 sec. Total RNA was extracted using the TRIzol method (Invitrogen) according to the manufacturer’s protocol except that 1-Bromo-3-ChloroPropane (BCP) (Sigma-Aldrich) 1 M was used instead of chloroform. DNase treatment was performed during 30 min at 37°C following the manufacturer’s protocol (TURBO DNase, kit Invitrogen), except that RNasine 0.36 U/μL (Promega) was also added. For digestive tracts containing blood, total RNA was extracted with the QIAamp RNA blood mini kit (Qiagen) following the manufacturer’s protocol. The recommended DNase I treatment (Qiagen) was also performed. RNA concentration was then measured using a nanodrop. Complementary DNAs (cDNAs) were synthesized from 25 ng/μL of total RNA using M-MLV Reverse Transcriptase (Invitrogen) or water (for negative RT) following the manufacturer’s protocol. Standard cDNAs were produced from 25 ng/μL of total RNA from whole females and were then diluted at 1:1000, 1:100, 1:10 and 1:1. All cDNAs were aliquoted and stored at −20°C until qPCR.

### qPCR

qPCR assays were performed with the Fast SYBR Green Master Mix method (Thermo Fisher Scientific) according to the manufacturer’s protocol and using specific primers (Sigma-Aldrich) for genes of interest (Table S1). Real-time qPCRs were run on an ABI 7500 Fast RT PCR machine and results were analysed with the 7500 Software v2.0.6. To quantify immune gene expression, data were analysed using the relative standard curve method. The average value of technical triplicates was normalized to the *S7* ribosomal average value for each sample and each gene. All qPCR within one experiment and independent replicates were performed using the same standard cDNAs batch. To quantify ZIKV RNA levels, qPCR assays were run with the comparative Ct (cycle threshold) method using *S7 ribosomal protein* gene as a standard gene for normalisation and according to the Taylor method (Taylor, Nadeau et al. 2019) to obtain a geomean of RQ=1 for the control group (no antibiotics-no sucrose) and relative RQ values for every other sample. To measure *16S* gene levels, 10 ng/μL of gDNA were used. qPCR assays were run with the comparative Ct method using *S7 ribosomal protein* gene as a standard gene for normalisation and according to the Taylor method (Taylor, Nadeau et al. 2019) to get a geomean of RQ=1 for the control groups (no antibiotics) and relative RQ values for every other sample.

### Statistics

Statistic results were obtained using a statistical software package (GraphPad Prism 9). For time course experiments, a Mann-Whitney test was performed for time point comparisons and a Kruskal Wallis test to test for the effect of time per treatment. Models including interactions (experiment X sugar X bacteria) were analysed with type-III ANOVA, whereas models without interactions (experiment X sugar or sugar X bacteria) were analysed with type-II ANOVA. Interactions with the experiment term were removed from the model as they were not statistically significant (p > 0.05). The p values indicate statistical significance of the treatment assessed with an ANOVA accounting for the experiment effect if there was one. Post hoc Fisher’s multiple comparisons were performed to get statistical significance of the desired pair-wise comparisons. For relative mRNA or gene detection using the comparative Ct method, Log2-transformed values of RQ values were used for statistical analyses. Statistical analyses of the infection prevalence were performed using a Chi-square test using the proportion of infected samples and the number of total samples per analysed group.

## Acknowledgements

We are grateful to Dr J.P. Parvy for helpful discussions and critical comments on the manuscript. We thank Dr A.B. Failloux (Institut Pasteur, France) for the Paea strain gift, Profs. M. Bouloy, R. M. Elliott and R. E. Randall for cell lines, J. Royle for ZIKV stock, and Dr C. Donald for ZIKV primers. This study was funded by the UK Medical Research Council (MC_UU_12014.8 to EP and AK).

## Author contributions

F.A.: Conceptualization, Data curation, Formal Analysis, Investigation, Methodology, Validation, Visualization, Writing – original draft, Writing – review & editing.

Sa.T.: Investigation, Project administration.

M.M.: Investigation, Writing – review & editing.

A.M.S.: Investigation, Resources, Writing – review & editing.

Se.T: Investigation, Writing – review & editing.

M.V.: Resources, Writing – review & editing.

A.M.: Investigation.

A.K.: Funding acquisition, Writing – review & editing.

E.P.: Conceptualization, Data curation, Formal Analysis, Investigation, Funding acquisition, Methodology, Project administration, Supervision, Validation, Visualization, Writing – original draft, Writing – review & editing.

## Conflicts of interest

The authors declare no conflict of interest.

## Supplementary material

**Figure S1.**
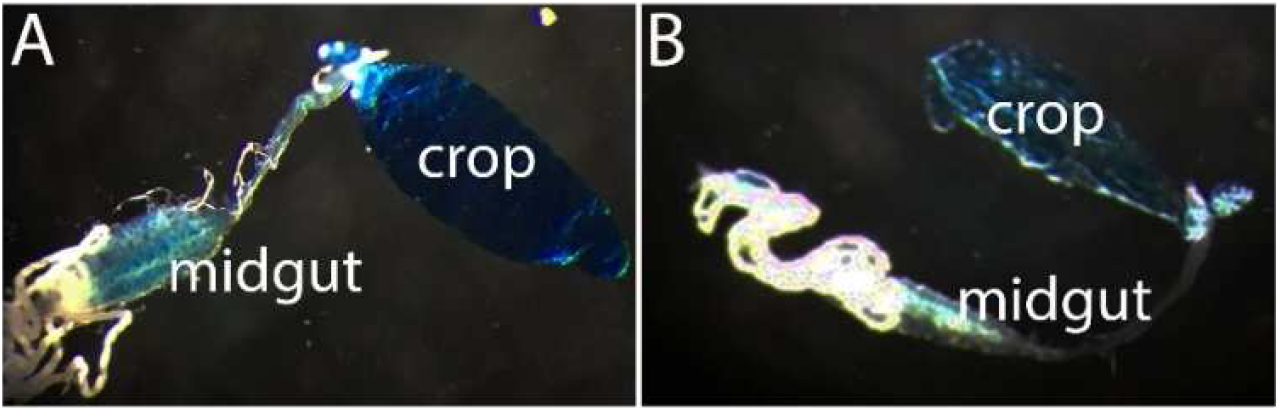
Sucrose solution is stored in the crop and intermittently relocated to the midgut. Pictures of digestive tracts dissected from females fed with a blue-stained 10% sucrose solution. (A) Picture of a representative gut minutes after sugar feeding showing sugar stored in the crop and relocated to midgut. (B) Picture of a representative gut one day after sugar feeding, showing sugar in the crop (although less than just after feeding) and little to no sugar solution in the midgut.

**Figure S2.**
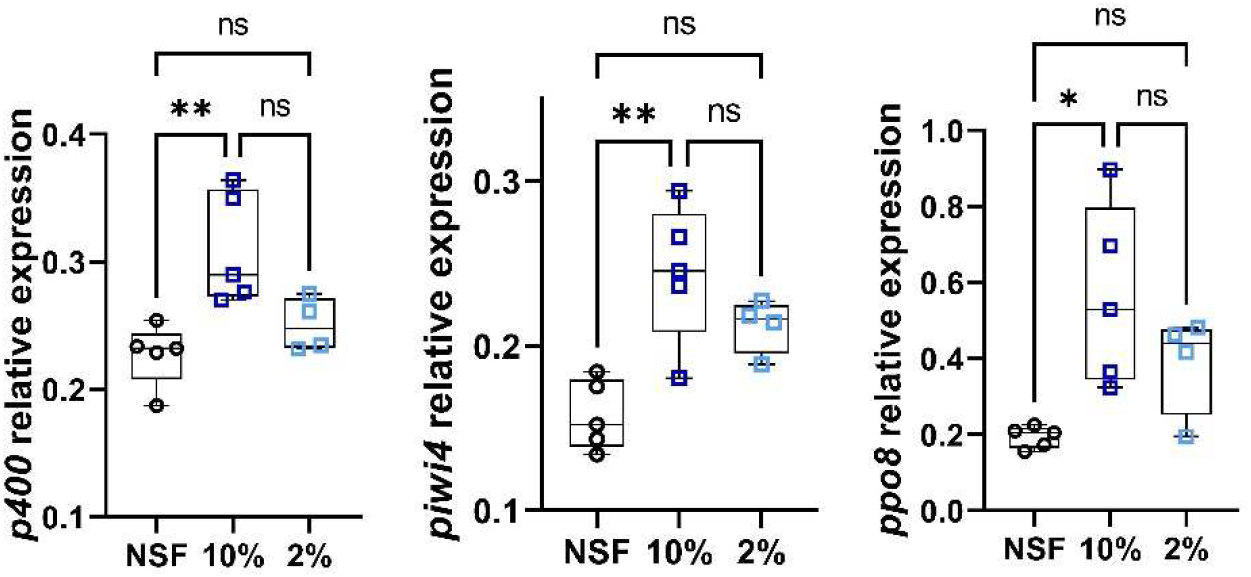
Sucrose increases immune gene expression in the digestive tract of *Ae. aegypti* females. Females that did not have access to sucrose for 48 h were either not fed (NSF), fed with 10% sucrose or 2% sucrose. Digestive tracts were dissected 16 h post sucrose feeding time. RNA transcript levels of *p400*, *piwi4* and *ppo8* were analysed by RT-qPCR. Box plots display the minimum, first quartile, median, third quartile, and maximum relative expression levels. Data were analysed by Kruskal-Wallis test followed by a Tukey’s multiple comparison test. N = 5 pools of 5 digestive tracts per condition. *, p value < 0.05; **, p value <0.01.

**Figure S3.**
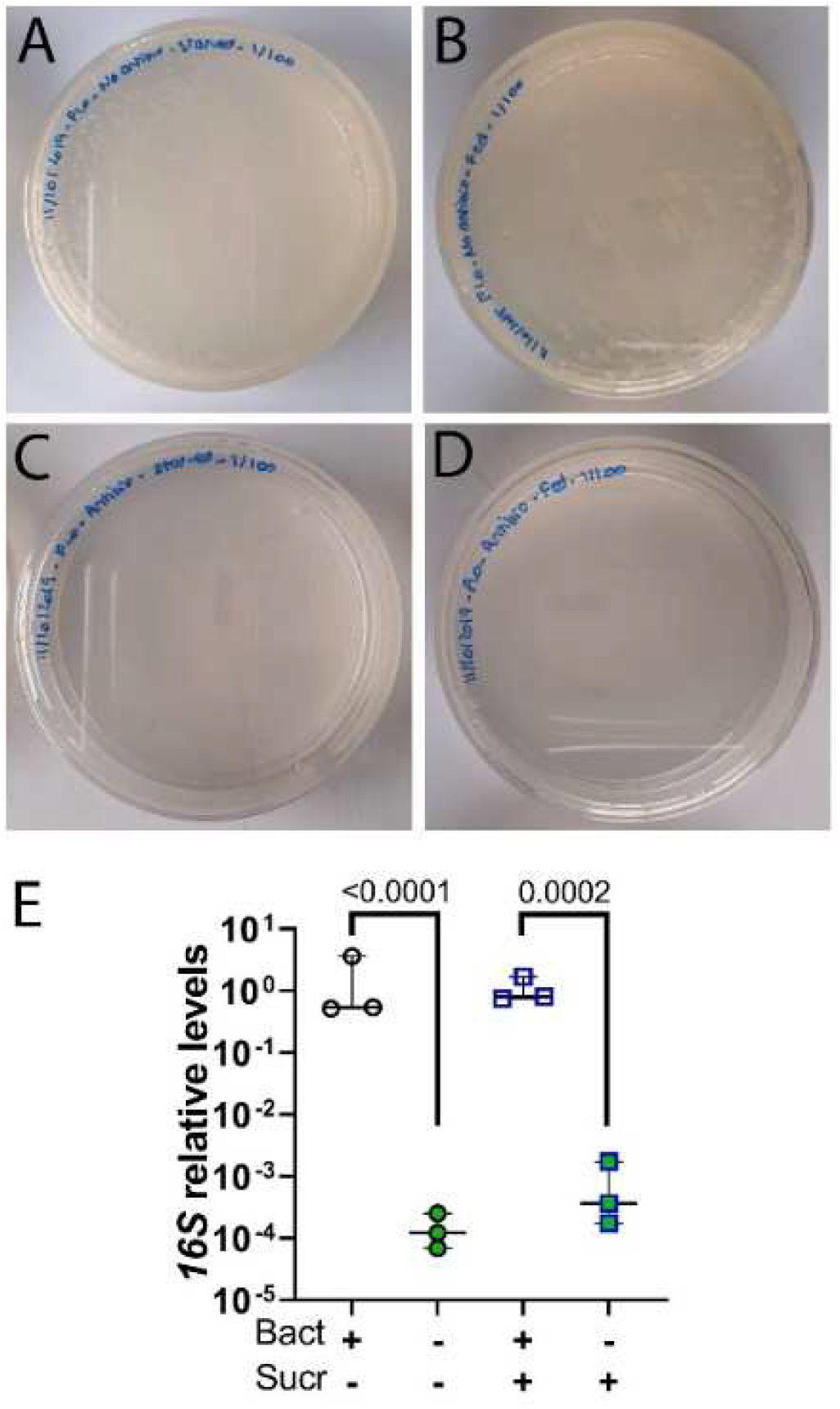
Validation of antibiotic treatment efficacy. Females, previously treated or not with antibiotics, did not have access to sucrose for 48 h to empty their crops and were then either not fed or fed with 10% sucrose. Digestive tracts were dissected (A to D) 16 h or (E) 18 h post sugar feeding time (E). (A to D) LB agar plates after plating homogenised and diluted (1/100) digestive tracts from (A) non fed females and (B) sucrose fed females not treated with antibiotics and digestive tracts from (C) non-fed females and (D) sucrose fed females treated with antibiotics. (E) *16S* relative gene levels in digestive tracts from the four populations, relative to *S7 ribosomal protein* gene, were analysed by qPCR. RQ for each sample was obtained as described [1], normalised to the *S7* ribosomal gene and as relative values to that of the respective control group (not treated with antibiotics, RQ geomean set to 1). Box plots display the RQ minimum, first quartile, median, third quartile, and maximum. Log2-transformed RQ values were analysed by ANOVA on matched values followed by a Holm-Sidak’s multiple comparison test. Bacteria in the digestive tracts of antibiotic-treated females were significantly depleted. Samples were similarly depleted of bacteria (No sucrose vs Sucrose: ns). N = 3 pools (from 3 independent experiments) of 5 digestive tracts per condition.

**Figure S4.**
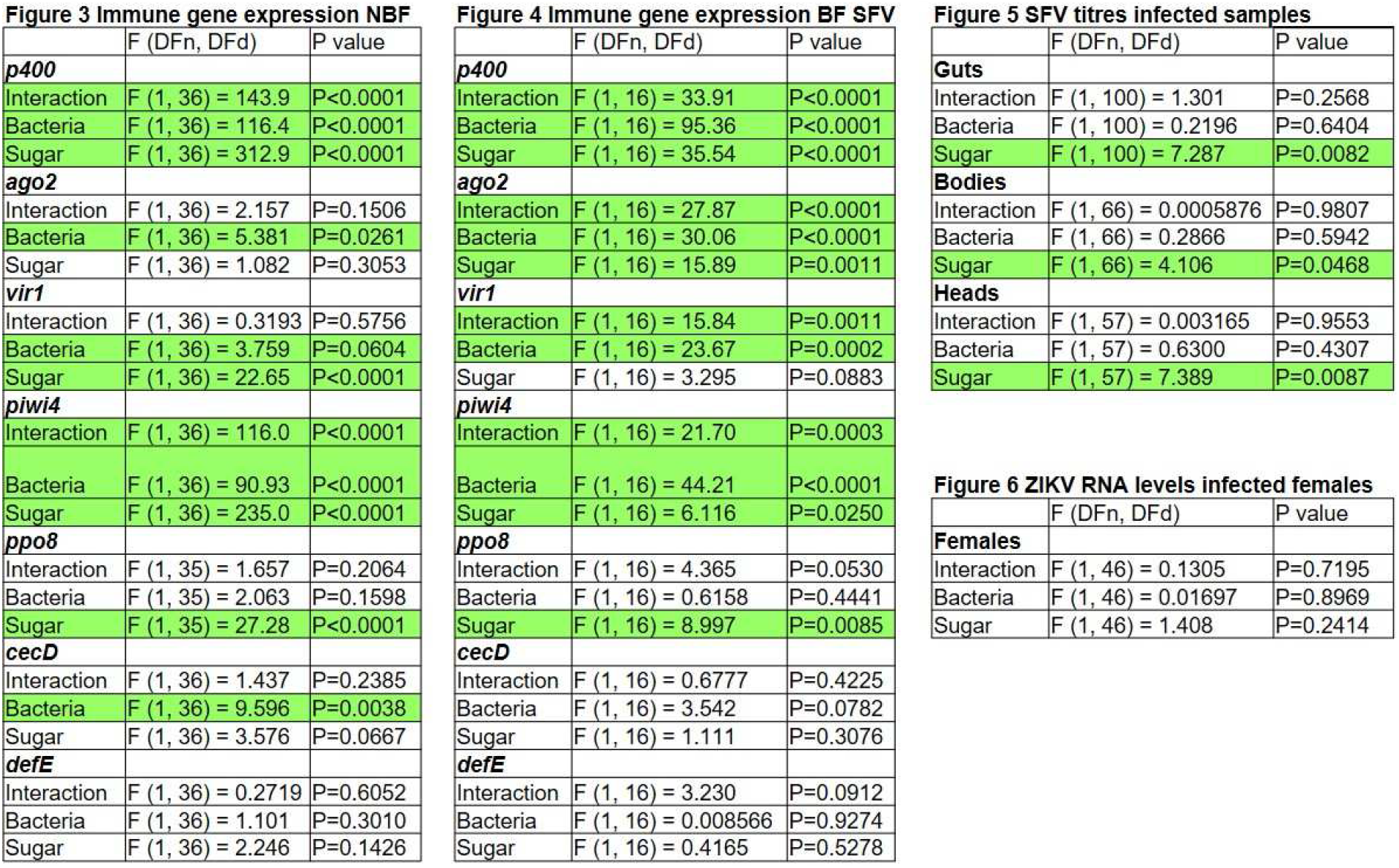
Two-way ANOVA statistical significance of treatments and interaction between treatments. Treatments: Sugar and Antibiotic (Bacteria) treatments. Significant treatment effects and interaction between treatments (p values < 0.05) are highlighted in blue.

**Table S1.**
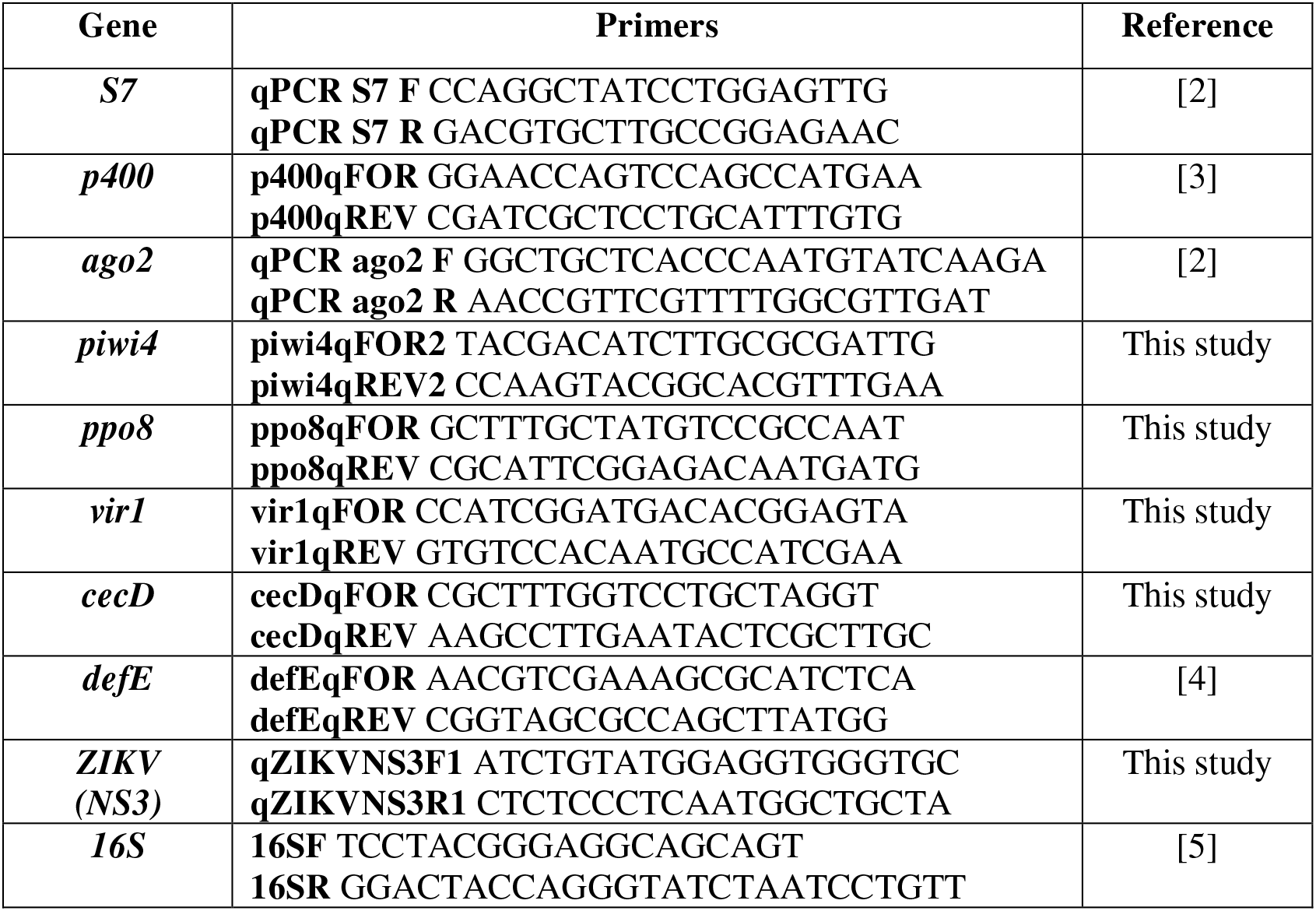
List of qPCR primers used in this study.

## Notes

### Competing Interest Statement

The authors have declared no competing interest.

